# Ancient DNA reveals the chronology of walrus ivory trade from Norse Greenland

**DOI:** 10.1101/289165

**Authors:** Bastiaan Star, James H. Barrett, Agata T. Gondek, Sanne Boessenkool

## Abstract

The search for walruses as a source of ivory –a popular material for making luxury art objects in medieval Europe– played a key role in the historic Scandinavian expansion throughout the Arctic region. Most notably, the colonization, peak and collapse of the medieval Norse colony of Greenland have all been attributed to the proto-globalization of ivory trade. Nevertheless, no studies have directly traced European ivory back to distinct populations of walrus in the Arctic. This limits our understanding of how ivory trade impacted the sustainability of northern societies and the ecology of the species they relied on. Here, we compare the mitogenomes of 27 archaeological walrus specimens from Europe and Greenland (most dated between 900 and 1400 CE) and 10 specimens from Svalbard (dated to the 18^th^ and 19th centuries CE) to partial mitochondrial (MT) data of over 300 modern walruses. We discover two monophyletic mitochondrial clades, one of which is exclusively found in walrus populations of western Greenland and the Canadian Arctic. Investigating the chronology of these clades in our European archaeological remains, we identify a significant shift in resource use from predominantly eastern sources towards a near exclusive representation of walruses from western Greenland. These results provide empirical evidence for the economic importance of walrus for the Norse Greenland settlements and the integration of this remote, western Arctic resource into a medieval pan-European trade network.

## Introduction

Atlantic walrus (*Odobenus rosmarus rosmarus*) ivory was a popular material for the manufacture of luxury art objects in medieval Europe. With isolated earlier and later exceptions, its use started in the 10^th^ century, peaked with the Romanesque art style of the 12^th^ century, and subsequently declined (1–4). This European demand for ivory has been considered a major economic incentive for exploration of the North Atlantic region and the Arctic (5–9). While the Atlantic walrus is widely distributed –with populations being found from Siberia to Canada (10)–, of all potential sources it is *Norse Greenland* for which ivory trade has been considered particularly important. First, the initial exploration and settlement of Greenland c.980-990 CE has been attributed to the hunt for ivory (7). Second, the 13^th^- and early 14^th^-century peak in transAtlantic trade to Greenland, and architectural (particularly church) investments there, have been similarly connected with the walrus (6, 9, 11–14). Finally, the abandonment of the Norse colony – its Western Settlement in the 14^th^ century and its (more southern) Eastern Settlement in the 15^th^ century – has been blamed on the declining popularity of walrus ivory in Europe and/or on a switch to alternative sources such as Svalbard or Russia (7, 15). To evaluate these hypotheses, we identify the Arctic sources of walrus imports to Europe between the 10^th^ and 15^th^ centuries using ancient DNA (aDNA).

The hunting of walruses for ivory by Norse Greenlanders is testified by archaeological finds of maxillae, maxillae fragments, tusk offcuts, postcanine teeth and carved objects (9, 16, 17). Post-cranial walrus bones (that represent hunting for meat rather than ivory) are rare from Norse sites in Greenland, particularly from the Eastern Settlement. Based on historical sources, most hunting took place further north along the west coast, mainly around Disko Bay (9, 18–21). The actual transport of walrus products –ivory, hide ropes and even a decorated walrus skull– from Greenland to Europe as gifts, tithes and trade goods is also described in historical sources. Although these sources vary in their historicity and are dated to the 13^th^ century CE or later, they describe practices thought (by their medieval authors and modern scholars) to have had an earlier origin (6, 7, 13, 21). Yet another major source of ivory was the Barents Sea region of Arctic Fennoscandia and Russia. This eastern source was documented as early as the late 9^th^ century CE, when the Arctic Norwegian chieftain Ohthere visited the court of King Alfred of Wessex in England (22, 23). The continued importance of this source is implied by the great abundance of walrus ivory known from medieval Novgorod –an important trading town with an extensive network into Arctic Fennoscandia and Russia (24, 25)– and by the hunting of the Arctic European walrus well into the 20^th^ century (26). Finally, walruses were also initially hunted in Iceland during its colonization in the late 9^th^ and 10^th^ centuries (7, 9, 27). By the 12^th^- 13^th^ centuries, when the island’s earliest laws and narrative texts were first recorded, Icelandic walruses were reduced to isolated visitors only (9, 28). Icelandic ivory finds were probably from local hunting during the Viking Age (7), but there exists no empirical evidence on the geographic origin of the ivory imported to European trading centers such as Trondheim, Bergen, Oslo, Dublin, London, Sigtuna and Schleswig during the chronology of the Norse Greenland settlements. Here we fill this gap in present knowledge.

Genetic analyses of modern walruses reveal significant population structure based on microsatellite variation (29–32), mitochondrial (MT) restriction fragment length polymorphism (RFLP) (29, 30) and partial MT sequence variation (33), which agrees with high levels of observed site fidelity (34, 35). Of particular relevance for this study, is the observation of a unique set of MT haplotypes that occur solely in western Greenland and the Canadian Arctic (29, 30, 33, 36, 37). We therefore presumed it possible to use mitogenomic data to identify the origin of archaeological walrus specimens traded from or via Greenland. Although aDNA can trace such specimens towards their biological source (38–40), it is difficult to apply this destructive technique to objects of fine art such as worked ivory. Nonetheless, partial walrus skulls (rostrums) with in-situ tusks were *also* transported to Europe (41). In rare instances these ‘ivory packages’ remain intact, while in others they were broken up to extract the ivory (41–46). These rostrums and rostrum fragments serve as proxies for the ivory they carried and can be sampled for biomolecular analyses without the need to damage ivory artefacts.

We analyze the mitogenomes of 24 archaeological walrus rostrums and three tusk offcuts. Most specimens have been dated by archaeological context, associated artefacts or (in one case) a runic inscription to the period of the Norse occupation of Greenland. Two late outliers that post-date this period are included for comparison, one from a context dating 1500- 1532 CE and one post-dating 1600 CE. Two other samples are undated, being from old excavations, but are probably medieval in origin. The specimens were originally obtained from excavations and subsequently archived in museum collections, although in Le Mans a rostrum with a 13^th^-14^th^ century runic inscription on one of its tusks has come down to posterity intact (41, 47). Four of our medieval rostrums serve as controls, having been excavated at the site of Igaliku (Gardar) in Greenland’s former Eastern Settlement (48). In addition, we analyze 10 control samples from 18^th^ and 19^th^-century Svalbard. We compare our data to 306 modern and 19^th^-century walrus specimens from the Barents Sea region, Greenland and the Canadian Arctic (32, 33, 36, 37) to infer the geographical origin of European walrus imports during the chronology of the Norse occupation of Greenland.

## Results

We obtained 520 million paired sequencing reads for 37 samples that passed downstream filtering (Fig. 1A, Table S1). The libraries of these samples contained 0.01 to 71% endogenous DNA and yielded 4.8 to 438-fold MT coverage (Table S2). The reads show the typical patterns of fragmentation and deamination expected from post-mortem degradation (Fig. S1). Including a *de novo* Pacific walrus MT sequence as outgroup (Supporting Information), a Bayesian phylogenetic analysis of 346 SNPs reveals two fully supported monophyletic clades in the archaeological Atlantic walrus data (Fig. 1B). The medieval walruses belong to either clade, whereas all the 18^th^ and 19^th^ century CE specimens from Svalbard fall into one clade. We tentatively define the clade containing the Svalbard samples as *eastern*, and the other clade as *western* (Fig. 1B). Of the four Greenland samples, two specimens fall into the western and two into the eastern clade (Fig. 1B). The two major lineages form distinct haplotype genealogies, with each clade separated by 21 or 18 substitutions from the most recent common ancestor (MRCA; Fig. S2). Using two different substitution rates, the time to the MRCA was estimated to have been 23,400 (95% HPD: 14,539-34,522) to 251,120 (95% HPD: 163,819-355,131) years ago.

**Fig. 1.**
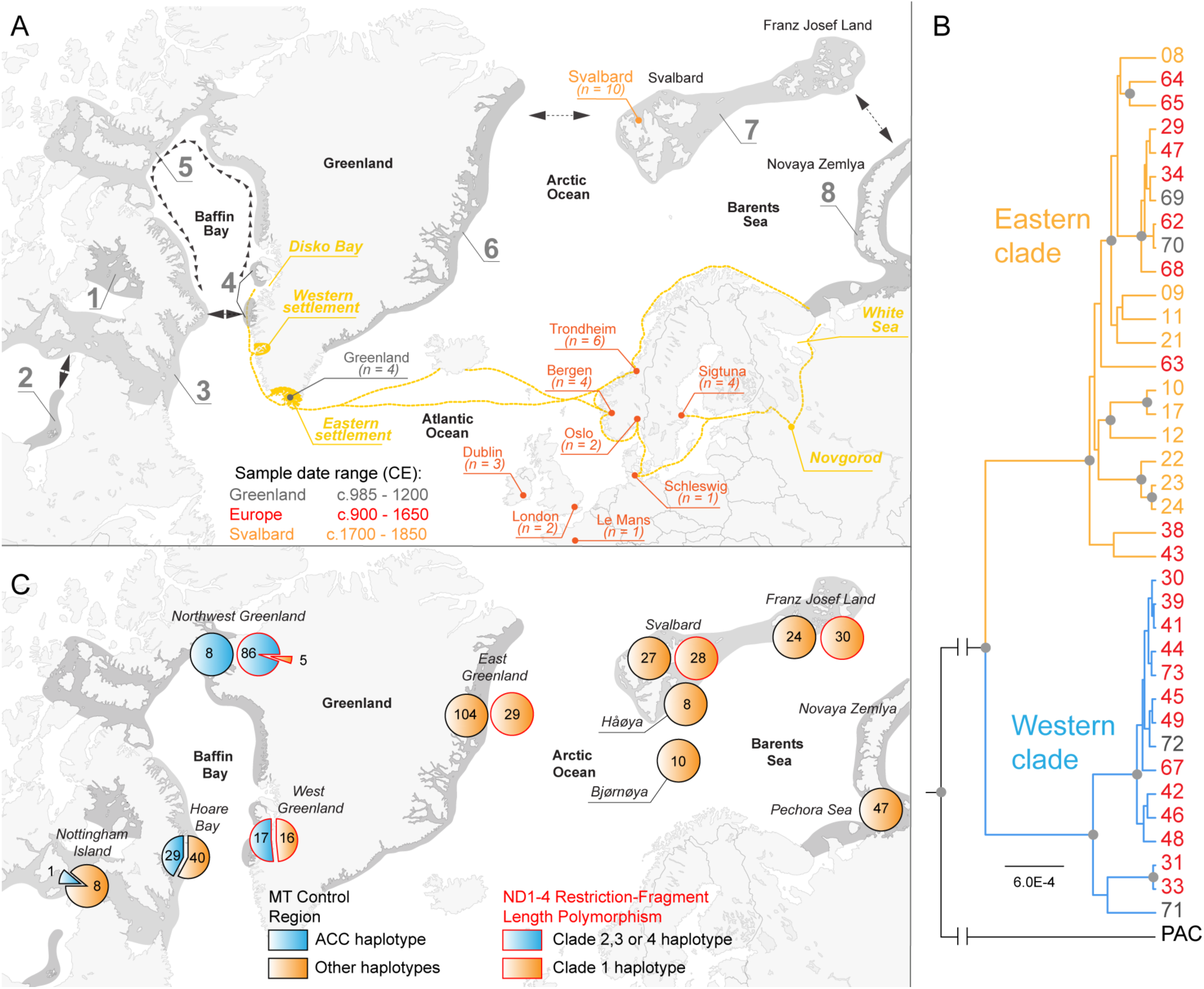
*(A)* Population distribution, historic trade routes and sample locations of Atlantic walrus in the northern Atlantic region. The range of modern Atlantic walrus (*dark grey*) and putative dispersal routes (*black arrows*) follow (58) and (31). Eight breeding populations are recognized (58); 1 - Foxe Basin, 2 - Hudson Bay, 3 - Hudson Strait, 4, - West Greenland, 5 - North Water, 6 - East Greenland, 7 - Svalbard/Franz Josef land, 8 - Novaya Zemlya. Historic trade routes from Greenland –including the location of Norse settlements– and northern Fennoscandia/Russia (*yellow*) indicate possible sources from which walrus ivory was exported to Europe during the Middle Ages. The Svalbard specimens (*orange*) were originally from hunting stations of the 1700s and 1800s. The other Atlantic walrus specimens (*red, grey*) were obtained from museum collections. *(B)* Bayesian phylogenetic tree obtained using BEAST (84) based on 346 mitochondrial SNPs using Pacific walrus (PAC) as an outgroup. Numbers represent the different specimens as listed in Table S1, and colors match the sampling locations as in Fig. 1A. Branches with a posterior probability of one (grey circles) are indicated. *(C)* Distribution of RFLP and control region (CR) haplotypes of modern Atlantic walrus populations. The RFLP clade classification follows Born, Andersen *et al.* (2001). The distribution of a distinct ACC CR haplotype is from 306 modern specimens (see material and methods).

A Principle Component Analysis (PCA) shows two significantly differentiated clusters supported by clade-specific SNPs that are located throughout the entire mitogenome (Fig. S3). We investigated if these clade-specific SNPs can be associated with RFLP (29, 30) and control region (CR) population data (32, 33, 36, 37). First, the RFLP studies focused on ND1, ND2, and ND3/4 MT gene regions (29, 30). We find 5 clade-specific SNPs in ND1, 3 in ND2 and 9 in ND3/4 (Supporting Information, Fig. S3C). At least one of these SNPs alters a restriction enzyme sequence motive depending on clade membership in each of these ND genes (Supporting Information). The combination of enzymes (29, 30) can therefore separate the western and eastern clades we identify in our mitogenomes –on multiple restriction sites and within each ND gene region. While these RFLP analyses report four MT clades, the data show a pronounced bimodal divergence, with the majority of distinctive restriction sites and highest statistical support observed between clade 1 and any combination of clades 2-4 (29, 30). Following Born *et al*. (2001), clades 2-4 comprise 94% of the Northwest Greenland and 52% of the West Greenland population (Fig. 1C). Crucially, these clade 2-4 haplotypes are exclusively identified in western Greenland, and are absent in the Northeast Atlantic (29, 30). Clade 1 is found in every individual in the Northeast Atlantic and in various proportions in western Greenland (Fig. 1C).

Second, we investigate a 499 base pair (bp) section (between 15328 and 15827 bp) of an extensive CR population dataset (32, 33, 36, 37). We observe three SNPs in this section, for which 12 out of 15 archaeological western clade specimens share a distinct A15564 C15760 C15779 haplotype (Supporting Information, Fig. S3C). Reanalyzing the CR data for this ACC haplotype (32, 33, 36, 37) we identify 38 out of 306 Atlantic walruses with the same haplotype (Supporting Information, Table S3). The ACC haplotype is fixed in the Northwest Greenland population, co-occurs mixed with other haplotypes in Canada, yet is absent from the Northeast Atlantic (Fig. 1C). The distribution of this CR haplotype is therefore analogous to the RFLP distribution. We derive that the RFLP and CR analyses detect variation explained by the monophyletic lineages of our medieval and 18^th^ and 19^th^ century CE specimens. Moreover, these analyses reveal a consistent distribution, whereby a subset of haplotypes is geographically restricted to western Greenland and Canada. Since the ACC haplotype of our western clade is only found in western Greenland and Canada, we conclude that the European archaeological specimens belonging to this clade came from Norse Greenland, either because they were hunted by the Norse or because they were traded from further north and west via contact with indigenous Dorset or Thule peoples (18, 19, 21, 49, 50). In contrast, haplotypes of the eastern clade may originate from either side of the Atlantic Ocean (Fig. 1C).

The dates of the archaeological walrus specimens cover the entire chronology of the Norse Greenland occupation, with two outliers post-dating its abandonment (Fig. 2A, Table S3). Before the founding of Greenland’s bishopric in the 1120s (probably by 1125) CE (51) – during the settlement phase and the early period of Norse occupation– we identify one western and six eastern clade specimens in Europe (Fig. 2A). In contrast, during the later occupation, between the 1120s and the end of the 14^th^ century, we observe 10 western and two eastern clade specimens. The timing of this significant increase (*p*-value = 0.0063, Fisher’s exact test) in western clade specimens thus coincides with new ecclesiastical infrastructure in Greenland (51) and the height of Romanesque ivory carving in Europe (3). We have not yet discovered European walrus rostrums that date specifically to the century of Norse Greenland’s abandonment (the 1400s). By this date walrus ivory had long gone out of fashion, with the Gothic style using different materials such as elephant ivory (4). Of the two later outliers in our dataset (both from Trondheim), one from the archbishop’s palace is of the western clade, dated 1500-1532. It may represent redeposition of an earlier find, or the latest known export of a walrus rostrum from Greenland. The second outlier postdates 1600 CE and is of the eastern clade.

**Fig. 2.**
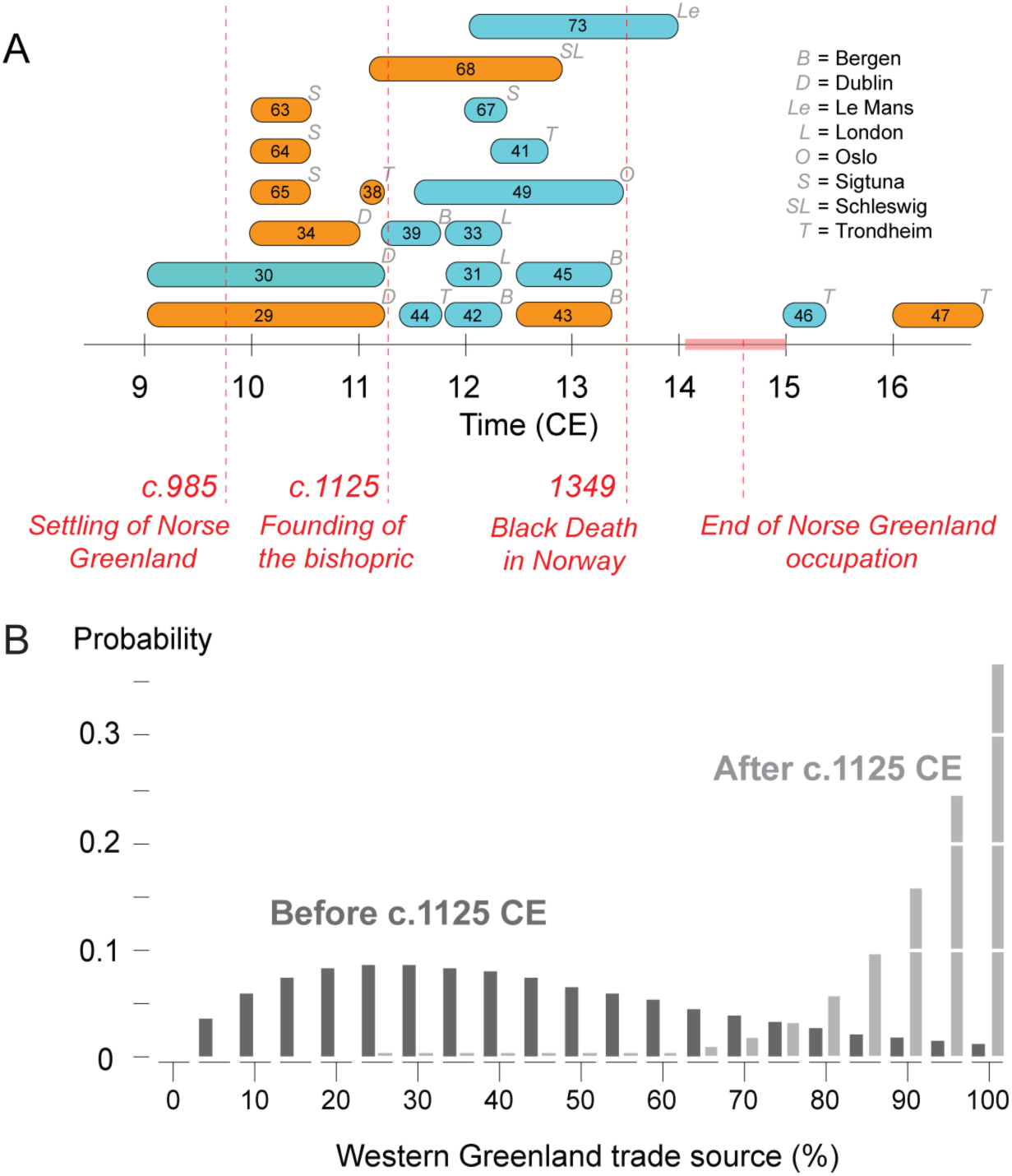
Chronology of Atlantic walrus specimens in Europe. *(A)* Archaeological Atlantic walrus specimens classified as western clade (blue) or eastern clade (orange) are plotted according to their estimated age range. The start and end of the Norse Greenland occupation, the founding of the bishopric and the arrival of the Black Death (plague) in Norway are indicated (*dashed red lines*). For each specimen, its location (light grey) is indicated. *(B)* Probability of obtaining the observed sample of eastern and western clade archaeological specimens as a function of a variable contribution of a western Greenland source. The probability was calculated for those samples obtained before (dark grey) or after (light grey) the founding of the Greenland bishopric, excluding the 16^th^ and 17^th^ century CE specimens.

Based on RFLP data, eastern clade specimens make up nearly half (48%) of the modern West Greenland population –which is closest to the Disko Bay hunting grounds of the Greenland Norse (Fig. 1C). Albeit a small sample size, the medieval Greenland specimens from Gardar also have such a 50/50 distribution, suggesting long-term temporal stability of these haplotype frequencies (Fig. 1B) and showing that eastern clade trade specimens could also have originated from Norse Greenland. We calculated the binomial probability of the observed ratio of western and eastern clade archaeological samples in the period *before* and *after* the founding of the Greenlandic bishopric c.1125 CE (excluding the two outliers from Trondheim), given variable contributions from western Greenlandic/Canadian and Northeast Atlantic/European Arctic sources. This probability is calculated assuming that populations in the Northeast Atlantic are fixed for the eastern clade, whereas those in western Greenland and Canada comprised a mixture of clades, following the RFLP frequencies (48% eastern clade) of the West Greenland population. The probability distribution of the samples dated before the bishopric’s founding shows evidence of geographical admixture, with a most likely contribution of a western source between 20 to 30% (Fig. 2B). In contrast, the samples dated after the founding and before c.1400 CE show evidence for a near 100% western clade source (Fig. 2B). In other words, in its early period, it is statistically unlikely that Norse Greenland was an exclusive source of walrus for Europe. Arctic Europe (the Barents Sea region) is the most likely alternative based on the Ohthere account of the late 9^th^ century. Iceland and (less likely, due to difficult summer ice conditions) Northeast Greenland are other possibilities (7, 21). In the later period between c.1125 and c.1400 CE, however, the number of observed eastern clade samples can –with high statistical probability– have come entirely from Norse Greenland together with the western clade specimens.

## Discussion

We reconstruct a chronology of long-distance ivory trade during the medieval period by investigating complete mitogenomes of archaeological Atlantic walrus specimens from Greenland, Svalbard and Europe. Specifically, we distinguish whether individual walruses were obtained from a western Greenland source and our work has evolutionary, ecological and archaeological implications.

We discover that Atlantic walrus comprises two major, monophyletic MT lineages. Several observations support a hypothesis that these lineages have evolved in glacial refugia on either side of the Atlantic Ocean. First, we estimate a divergence date between 23400 and 251120 years ago, well before or during the last glacial maximum (LGM). Second, (sub)fossil walrus bones dated over 10k years BP have been found on both the European and the North American continent *at lower latitudes* compared to the walrus’s modern distribution (52, 53). Third, glacial refugia on either side of the Atlantic have also been proposed to explain (partial) MT divergence in marine mammals with a similar coastal ecology like walrus, such as harbor seal (*Phoca vitulina*) (54), harbor porpoise (*Phocoena phocoena*) (55) and grey seal (*Halichoerus grypus*) (56). Based on these observations, we conclude that the two MT lineages have evolved in spatial separation on either side of the Atlantic Ocean.

Such a scenario further explains the distinct geographical distribution of RFLP and CR population data, which show that haplotypes associated with the western clade occur solely in western Greenland and Canada (29–32, 37). The lack of such haplotypes in the Northeast Atlantic, despite the genetic analysis of hundreds of specimens covering multiple decades (29–32, 37), indicates that gene flow has been asymmetrical, with eastern clade females dispersing to the western Atlantic but not *vice versa*. This dispersal follows the direction of the East Greenland Current (57), suggesting that ocean currents influence the tendency of walrus dispersal. Historically, a similar asymmetrical pattern has been observed on a regional scale in Baffin Bay –with walruses migrating counter-clock-wise, following the direction of the coastal currents and the breakup of sea ice (31, 58). Our data suggest that dispersal has been persistent in its direction since these two MT lineages came into secondary contact. As a result, eastern clade walruses are now found in the entire range of Atlantic walrus, yet western clade walruses have not yet been observed in the Northeast Atlantic.

With our discovery of two MT lineages we provide quantitative evidence on the geographic origin of walrus imports to medieval Europe, and with that re-evaluate past hypotheses on the role of walrus ivory in the origins, efflorescence and collapse of Norse Greenlandic society. Only one of our seven samples predating the 1120s CE is of the western clade. The other six eastern clade samples of this period could have come from a variety of sources, including Greenland, but it is probable that most derived from the Northeast Atlantic. Thus the theory that walrus ivory was a primary motive for the initial exploration and settlement of Greenland may need reconsideration (7). Conversely, 10 of 12 specimens dating between the 1120s and c.1400 CE are of the western clade and arrived in Europe via Norse Greenland. Moreover, the observed ratio of eastern and western clade specimens has a high probablility of deriving exclusively from western Greenland. Thus, the documented heyday of Norse Greenlandic settlement, trade and architectural elaboration (particularly evident in churches and their accoutrements) – between the 12^th^ and 14^th^ centuries – did coincide with exports of walrus ivory. In fact, our data suggest that the Greenland trade of this commodity may have held a near monopology in western Europe. The historically and/or archaeologically attested walrus hunts of the 9^th^-10^th^ centuries in the Barents Sea region and Iceland may have declined or ended by the early 12^th^ century, either because of rising Greenlandic exports or local reduction/extirpation of populations.

The reasons for the signficant shift in trade cannot be inferred from aDNA alone and historically contingent local factors, as well as broader socioeconomic and environmental developments can be invoked as potential contributors. In 13^th^-14^th^ century Iceland it was believed that the Greenlanders used walrus products as gifts to influence the policies of Scandinavian monarchs, for example in the story of Einar Sokkason (*Grænlendinga tháttr*) (2). Although the historicity of Einar’s use of walrus ivory to secure a Greenland bishop and episcopal see in the early 12^th^ century cannot be confirmed, tithes (including papal dues) were paid in this material during the 13^th^ and 14^th^ centuries (1, 6). Thus both local initiative and the reach of panEuropean church infrastructure potentially played a role in the naissance and maintenance of the Greenland ivory trade. Moreover, the 11^th^ to 13^th^ centuries represent a period of major demographic and economic growth in Europe, in part due to environmental conditions favourable to cereal agriculture (59). A growing urban and elite demand was served by transport from increasingly distant sources in a process of proto-globalization. Our discovery shows that Greenland was well integrated into this network.

Less can be said about the end of the Greenland colony based on the evidence here. Is the absence of sampled (or known) European finds of walrus rostrums from the 15^th^ century evidence for the end of trade? Is the single, 16^th^-century western clade specimen an example of the last, isolated, Greenland export, or is it a redeposition of an earlier find? These are classic challenges of archaeological interpretation. Nonetheless, it is a conspicuous observation that Greenland may have been the exclusive supplier of walrus ivory to Europe between the 1120s and the 14^th^ century. The demise of Norse Greenland would therefore have reduced European supplies of this raw material, whereas a decline in demand would have undermined Greenland’s social and economic organization. Whatever other factors have been influential –from the Little Ice Age (60–63), to gradual out-migration (62, 63), to the impact of the Black Death (1346–1353) on European markets (2, 59)– the cessation of trade in walrus ivory must have been significant for the end of Greenland’s Eastern and Western settlements.

## Conclusion

Here, we show that the Atlantic walrus comprises two monophyletic MT lineages of which one is exclusively found in western Greenland and Canada. By analyzing archaeological walrus remains in Europe, this discovery allows us to infer an increasing trade of walrus ivory from Norse Greenland. Greenland is often discussed as a general example of both human resilience and vulnerability in the face of environmental and economic change (61, 64–67). Thus, the implications of this study –that the influence of ecological globalization for the Greenlandic Norse started small yet became paramount– extend far beyond medieval Europe.

## Material and Methods

We sampled 37, morphologically identified walrus bone and ivory specimens from Western Europe, Greenland and Svalbard (Fig. 1A, Table S1). The Svalbard specimens were dated based on the documented use of the hunting stations from which they were collected. The Le Mans specimen was dated based on the characteristics of a runic inscription on an *in situ* tusk. The other specimens were dated by archaeological context and/or associated artefacts. Direct radiocarbon dating was not used because precise marine reservoir correction requires a ΔR value which cannot be predicted without catch location (68).

All DNA extraction and pre-PCR protocols were performed in a dedicated laboratory at the University of Oslo following strict aDNA precautions (69, 70). Samples were exposed to UV (Supporting Information) before milling using a custom designed stainless-steel mortar (71) or a Retsch MM400 mixer mill. Extraction used a combined bleach and pre-digestion protocol (72), apart from the Greenland samples for which only pre-digestion was used (73). Blunt-end Illumina libraries were built (39, 74) amplifying ligated DNA with sample-specific seven bp indexes in the P7 primer (Supporting Information). Libraries were sequenced (125 bp paired-end) on an Illumina HiSeq 2500 and demultiplexed allowing zero mismatches in the index tag.

Sequencing reads were processed using PALEOMIX (75) (collapsing forward and reverse reads using AdapterRemoval v1.5 (76)) and aligned to the Pacific walrus (*O. r. divergens*) nuclear genome (77) and the Atlantic walrus mitogenome (78) with BWA *aln* v.0.7.5a-r405 (79). Read data from the Pacific walrus genome project (77) were aligned to the Atlantic walrus mitogenome to obtain a Pacific walrus MT sequence (Supporting Information). Alignments with a quality score (MapQ) of <25 were removed and aDNA damage patterns were investigated using mapDamage v.2.0.6 (80). Genotypes were obtained using GATK v. 3.4.46 (81) *Haplotypecaller* with ploidy set to 1, after duplicate removal (Picard Tools v.1.96) and indel realignment (*GATKs IndelRealigner*). Genotypes were jointly called using default settings (GATKs *Genotypecaller*), and filtered with BCFTOOLS v. 1.6 (82) using filters -i ’FS<60.0 && SOR<4 && MQ>30.0 && QD > 2.0’ and --SnpGap 10. Indels were excluded using VCFTOOLS v.0.1.14 (83) and genotypes with a quality below 15 and a read depth below 3 were set as missing.

Phylogenetic analyses were performed using BEAST 2.4.7 (84) with Pacific walrus as outgroup. Following jModeltest 2 (v0.1.10) (85), the HKY model was implemented with the Yule tree prior and a strict clock model. Timing to the most recent common ancestor (MRCA) was estimated using a faster rate of 0.75 x 10^-7^ substitutions/site/year from southern elephant seal CR data (86) and a slower rate of 0.7 x 10^-8^ estimated from baleen whale cytochrome b data (87, 88), because the CR mutates relatively fast compared to other parts of the mitogenome (89). The MCMC (10 million gen, pre-burnin 1 million gen) was sampled every 1000 trees and ESS values above 200 confirmed convergence (Tracer v.1.6.0 (90)). The maximum clade credibility tree was drawn after a 10% burnin using TreeAnnotator and the tree was visualised in FigTree (v.1.4.3). A haplotype genealogy was drawn using Fitchi (91). Differentiation among lineages was assessed using smartPCA, EIGENSOFT v.6.1.4 (92) obtaining clade supporting SNPs using *snpweightoutname* (Fig. S3). Clade specific modification of RFLP motivs (29, 30) was obtained using *Find Motif*, IGV (93) (Supporting Information). CR population data (32, 33, 36, 37) were aligned to the Atlantic walrus MT genome and genotypes at position 15564, 15760 and 15779 were scored in IGV (94) (Supporting Information, Table S3). Finally, the binomial probability of observing different western and eastern clade ratios before and after c.1125 CE was calculated by simulating variable contributions of a western Greenlandic/Canadian and Northeast Atlantic source, assuming 100% eastern clade specimens in the Northeast Atlantic populations and 48% in western Greenland/Canada. Probabilities for source admixture were scaled to one for each period.

## Acknowledgements

This work was supported by the Nansenfondet, the Research Council of Norway projects 262777 and 230821 and the Leverhulme Trust project “Northern Journeys” MRF-2013-065. We thank M Skage, S Kollias, A Tooming-Klunderud and H Rydbeck from the Norwegian Sequencing Centre. Samples (and information regarding their chronology) were kindly provided by: National Museum of Denmark (J Arneborg), Natural History Museum of Denmark (KM Gregersen, KH Kjær), NTNU Museum of Natural History and Archaeology (A Christophersen, JA Risvaag, J Rosvold), University Museum, Bergen (G Hansen, AK Hufthammer, S Nordeide), Museum of Cultural History, University of Oslo (M Vedeler), NIKU (LM Fuglevik), National Museum of Ireland (M Sikora, A Halpin), MOLA (L Blackmore, A Pipe), Sigtuna Museum (A Söderberg), Schleswig-Holsteinische Landesmuseen Schloss Gottorf (V Hilberg, U Schmölcke) and Le musée Vert, Le Mans (N Morel). AH Pálsdóttír and S Wickler also helped obtain samples.

## Data availability

All ancient read data are available at the European Nucleotide Archive (ENA, www.ebi.ac.uk/ena) under study accession number PRJEB25536.

## Author contributions

BS, JHB and SB designed the study. AG performed the laboratory work. BS performed the genetic analyses. SB performed the Bayesian analyses. BS, JHB & SB interpreted the results. JHB identified and selected ancient samples. BS, JHB & SB wrote the paper.

## Supporting Information

### Supplementary Notes

#### 1. Generating a Pacific walrus (*Odobenus rosmarus divergence*) mitochondrial genome sequence

The publicly released genome sequence of the Pacific walrus (77) includes a mitogenome assembly from Atlantic walrus (78), Genbank ID NC_004029.2). Hence, no *de novo* Pacific walrus mitochondrial reference genome sequence is currently available. To create a Pacific walrus MT genome sequence to root our phylogenetic analyses, we obtained a subset of 169,056,390 paired reads generated for the Pacific walrus genome assembly project (file SRR575502, ENA project ID PRJNA167474, (77)) and aligned these to the Atlantic walrus MT reference genome with PALEOMIX (75) using BWA *mem* v.0.7.5a-r405 (79). Both collapsed and paired reads were used, resulting in a 13-fold coverage of the mitogenome. The Pacific walrus alignment data and the archaeological Atlantic walrus data were further processed simultaneously.

#### 2. DNA extraction and library preparation

All DNA extraction and pre-PCR protocols were performed in a dedicated laboratory at the University of Oslo following strict aDNA precautions (69, 70). Samples were exposed to UV (10 min) on each side (total dosage of 4800 J/m2) before cutting. Dust was removed with UVed milliQ and cut fragments were again exposed to UV (10 min) on each side (total dosage of 4800 J/m2) before milling using a custom designed stainless-steel mortar (71) or a Retsch MM400 mixer mill. Extraction used a combined bleach and pre-digestion protocol (72), apart from the Greenland samples for which only pre-digestion was used (73). Bleach washes were done in duplicate (150-200 mg of powder each) (72), washed with H_2_O and pre-digested, followed by an overnight, second digestion (73). The two eluates were combined and concentrated (Amicon-30kDA Centrifugal Filter Units), extracting DNA using Minelute (Qiagen) according to manufacturer’s instructions. DNA was eluted in 60 µl pre-heated (60°C) EB buffer, incubating for 15 min at 37°C (95). Negative controls were included in all extraction experiments. Blunt-end Illumina libraries were built (39, 74) amplifying ligated DNA with sample-specific seven bp indexes in the P7 primer. PCRs were done in triplicate, 25 µl reactions (1.25 U PfuTurbo Cx Hotstart DNA Polymerase (Agilent), 1x buffer, 0.2 mM per dNTP, 0.2 µM P7 index primer, 0.2 µM P5 IS4 primer and 0.4 mg/ml BSA) for 13 cycles (2 min at 95°C, 13 cycles of 30s at 95°C, 30s at 60°C and 70s at 72°C, final extension of 10 min at 72°C). Amplified products were cleaned using Agencourt® AMPure XP beads at a 1:1.7 ratio, eluted in 30 µl of molecular grade H_2_O, and quantified using a Bioanalyzer 2100 (Agilent). Libraries were sequenced (125 bp paired-end) on an Illumina Hiseq 2500 and demultiplexed allowing zero mismatches in the index tag.

#### 3. Assessing Restriction Fragment Length Polymorphism in Atlantic walrus

Significant population genetic structure based on mtDNA variation in Atlantic walrus has been detected using Restriction Fragment Length Polymorphism (RFLP) analysis targeting the ND1, ND2, and ND3/4 genes (29, 30). Specifically, several RFLP mtDNA haplotypes have been reported that are solely found in specimens obtained from western Greenland and the Canadian Arctic, and these haplotypes have therefore been suggested to be diagnostic markers, unique to these western populations (29, 30). We here assess whether the RFLP haplotypes obtained in these earlier studies are linked to the observed monophyletic divergence between the western and eastern clade of Atlantic walrus in our ancient samples. First, we expect diagnostic SNPs supporting the two clades found in the present study to be located in the ND1, ND2 and ND3/ND4 region. Moreover, at least some of these diagnostic SNPs should alter the sequence motive of restriction fragment binding sites specific to the restriction enzymes used in the RFLP studies.

In our mitogenomes, we identify five diagnostic SNPs with principle component weightings greater than 1.25 (smartPCA, EIGENSOFT v.6.1.4 (92)) in ND1 (between 2752 and 3708 bp), three SNPs in ND2 (between 3920 and 4963 bp) and nine SNPs in ND3/ND4 (between 9485 and 11568 bp). In each of these three regions, we find at least one SNP that alters the sequence motive of a restriction enzyme according to each monophyletic clade. In ND1 at position 2982, a G/A polymorphism distorts the GGCCC (western) binding motive of *Sau961* and *HaeIII* into AGCCC (eastern). In ND2 at position 4597, an A/G polymorphism alters the CGACT (eastern) motive restricted by *Hinf1* to CAACT (western). In ND3/ND4 at position 9658, a G/A polymorphism distorts the GGCC (western) binding motive of *HaeIII* into AGCC (eastern). Finally, in ND3/ND4 at position 10616, a T/C polymorphism distorts the TTTAAA (western) binding motive of *Dra1* to ATTAAA (eastern). The haplotype separation obtained by the earlier RFLP studies using these specific restriction enzymes can therefore be explained by diagnostic SNP differentiation that is directly linked to the two monophyletic clades discovered in our ancient mitogenomes.

#### 4. Atlantic walrus Control Region (CR) Polymorphism

Within 27 Atlantic walruses sampled from both sides of the Atlantic Ocean, significant population differentiation –identifying a group of monophyletic western Greenland haplotypes– has been detected based on the combined data from the ND1, COI and the mtDNA control region (CR) (33). Using data from the CR region alone, however, extensive population studies targeting hundreds of individuals show that this region lacks discriminatory power to confidently resolve Atlantic walrus populations within the Northeast Atlantic (32) nor distinguish between western and eastern Atlantic populations (36, 37). Nevertheless, several haplotypes were found only in the western Greenland region and not in any of the studied populations in East Greenland, Svalbard or Franz Josef land (36, 37). This observation suggests the existence of CR haplotypes that are only found in walruses in western Greenland. We here associate these CR haplotypes to the monophyletic divergence observed in our historic samples and the 27 modern specimens studied by Lindqvist *et al.* (33).

In our mitogenome data, we identify three SNPs (A/G_15564_, C/T_15760_ and C/T_15779_) with principle component weightings above 1.25 (smartPCA, EIGENSOFT v.6.1.4 (92)) that fall within the region of the CR covered by all earlier studies, between 15328 and 15827 bp (32, 33, 36, 37). In contrast to the diagnostic SNPs in the ND1, ND2, ND3 and ND4 region (see Supplementary Note 2), neither of these CR SNPs *alone* is fixed in either clade, reflecting the lack of power of the CR to differentiate the observed divergence over the entire mitogenome. Yet, we note that the A15564 C15760 C15779 haplotype occurs in 12 out of 15 of the archaeological specimens from the western clade while it is not found in the eastern clade.

We investigate the occurrence of this specific ACC haplotype in 105 publicly available CR sequences (Table S2) (32, 33, 36, 37) obtained from Genbank. These haplotypes represent over 300 individual Atlantic walruses sampled from multiple locations east and west of Greenland (Figure 2C). For walruses from East Greenland, Svalbard, Frans-Josef land and the Pechora Sea we report the specimen numbers of (32) as this study includes samples analysed by (33). Historic data from Håøya and Bjørnøya were obtained from (37) and the Northwest Greenland samples were obtained from (33). From (36) we selected only data from those two locations (Nottingham Island and Hoare Bay) from which more than two specimens were sampled. All obtained sequences were aligned to the Atlantic walrus reference genome using BWA *mem* v.0.7.5a-r405 (79), and their genotypes at position 15564, 15760 and 15779 were scored in IGV (94).

We identify 38 out of 306 modern Atlantic walruses with the same CR haplotype (Table S2). None of the modern samples obtained from populations in the Northeast Atlantic contained this ACC haplotype in the CR, several samples in Canada have this haplotype, all samples from Northwest Greenland have this haplotype (Table S2, Figure 1C). This result shows that the ACC haplotype we observe in our archaeological mitochondrial data is restricted to modern walrus populations in western Greenland and the Canadian Arctic.

## Supplementary tables

**Table S1.**
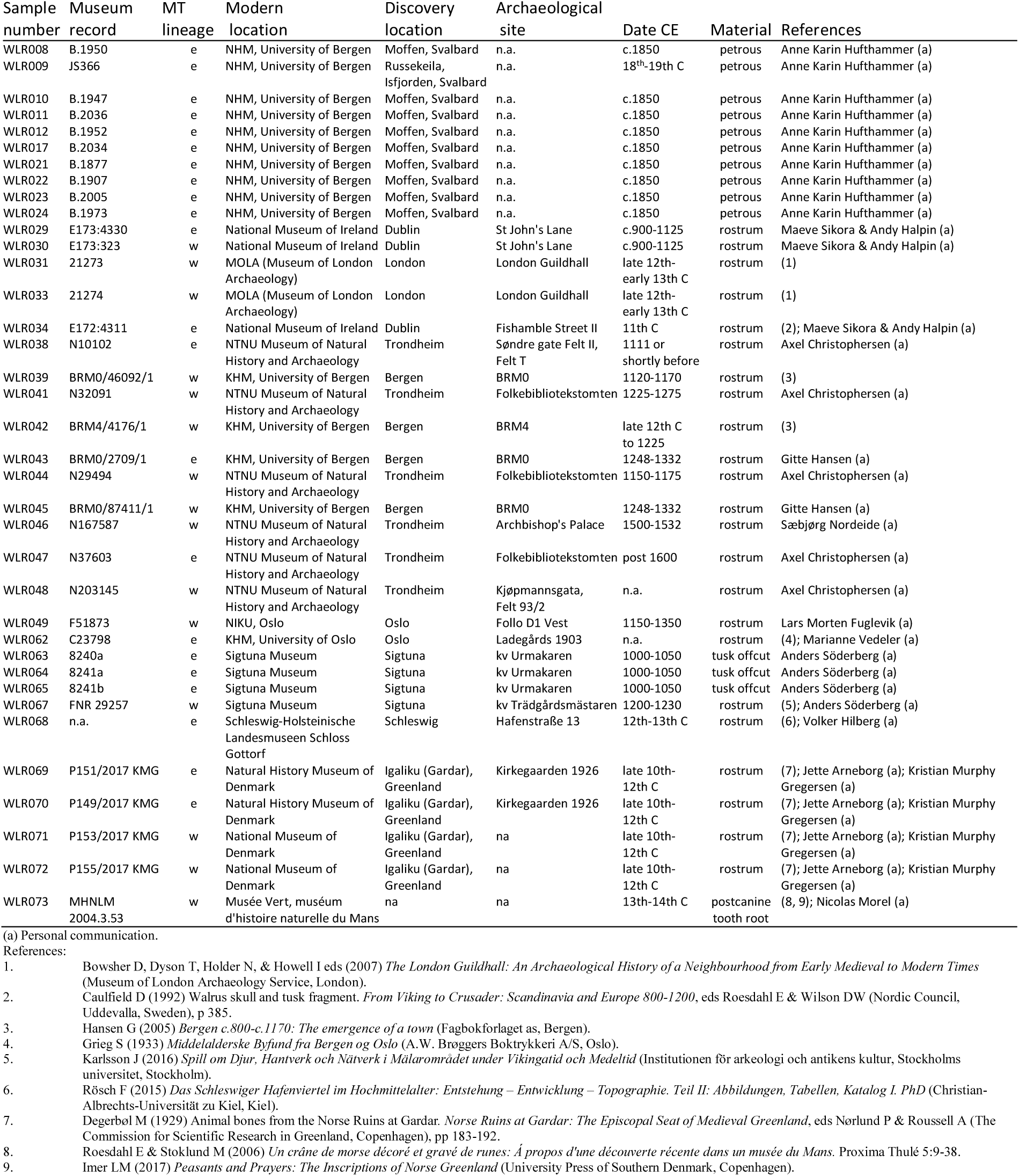
Sample details of archaeological Atlantic walrus specimens. The inferred genetic (*e; eastern clade, w; western clade*) MT lineage of each specimen is also indicated. Based on anatomical representation, animal size, archaeological context and/or MT haplotype all bones except two London specimens can be confidently interpreted as from separate animals. The two London specimens could be parts of the same walrus rostrum (representing left and right alveoli) although they were obtained from different archaeological layers. All samples were taken with museum permission and transported across international borders with the relevant CITES permits.

**Table S2.** Sequencing details of 37 archaeological Atlantic walrus samples. Estimates for library clonality and endogenous DNA content were obtained by aligning reads to the nuclear Pacific walrus reference genome (77). To obtain mitochondrial (MT) data, reads were aligned separately to the Atlantic walrus mitogenome (78).

| Sample ID | Reads<br>(millions) | Clonality (%) | Endogenous<br>DNA<br>(fraction) | MT fold<br>coverage | Insert<br>length(bp) |
| --- | --- | --- | --- | --- | --- |
| WLR008 | 13 | 0.01 | 0.44 | 43 | 70 |
| WLR009 | 16 | 0.01 | 0.09 | 17 | 80 |
| WLR010 | 19 | 0.01 | 0.33 | 59 | 79 |
| WLR011 | 6 | 0.00 | 0.23 | 8 | 62 |
| WLR012 | 5 | 0.00 | 0.43 | 30 | 68 |
| WLR017 | 20 | 0.01 | 0.68 | 125 | 78 |
| WLR021 | 15 | 0.01 | 0.71 | 80 | 91 |
| WLR022 | 15 | 0.01 | 0.67 | 89 | 87 |
| WLR023 | 12 | 0.01 | 0.35 | 73 | 96 |
| WLR024 | 16 | 0.01 | 0.69 | 49 | 81 |
| WLR029 | 12 | 0.02 | 0.29 | 52 | 69 |
| WLR030 | 17 | 0.03 | 0.17 | 74 | 72 |
| WLR031 | 9 | 0.02 | 0.16 | 81 | 72 |
| WLR033 | 17 | 0.03 | 0.01 | 8 | 94 |
| WLR034 | 30 | 0.42 | 0.02 | 7 | 58 |
| WLR038 | 5 | 0.04 | 0.35 | 16 | 76 |
| WLR039 | 8 | 0.03 | 0.01 | 13 | 61 |
| WLR041 | 1 | 0.01 | 0.48 | 10 | 70 |
| WLR042 | 11 | 0.02 | 0.01 | 13 | 74 |
| WLR043 | 13 | 0.08 | 0.00 | 9 | 71 |
| WLR044 | 10 | 0.01 | 0.49 | 14 | 80 |
| WLR045 | 12 | 0.02 | 0.11 | 24 | 75 |
| WLR046 | 10 | 0.03 | 0.03 | 22 | 97 |
| WLR047 | 11 | 0.05 | 0.30 | 9 | 64 |
| WLR048 | 11 | 0.03 | 0.42 | 179 | 79 |
| WLR049 | 12 | 0.02 | 0.02 | 9 | 79 |
| WLR062 | 15 | 0.06 | 0.49 | 295 | 62 |
| WLR063 | 12 | 0.01 | 0.06 | 46 | 77 |
| WLR064 | 9 | 0.01 | 0.03 | 14 | 72 |
| WLR065 | 12 | 0.02 | 0.37 | 438 | 73 |
| WLR067 | 6 | 0.01 | 0.08 | 13 | 69 |
| WLR068 | 10 | 0.02 | 0.19 | 104 | 67 |
| WLR069 | 23 | 0.01 | 0.01 | 5 | 76 |
| WLR070 | 27 | 0.01 | 0.02 | 10 | 72 |
| WLR071 | 27 | 0.01 | 0.01 | 6 | 69 |
| WLR072 | 35 | 0.01 | 0.02 | 10 | 69 |
| WLR073 | 16 | 0.01 | 0.15 | 68 | 72 |

**Table S3.** Genotypes for three control region SNPs in 105 Atlantic walrus control region sequences. These SNPs are located between 15328 and 15827 bp. For each of the sequences, we show their Genbank accession number, study from which the data was obtained, and whether the individuals with these sequences originated from western Greenland or Canada (West) or the Northeast Atlantic (East). The MT location and alleles (between brackets) for each SNP are given. The A_15564_ C_15760_ C_15779_ haplotype that only occurs in western Greenland and the Canadian Arctic is highlighted (**bold**).

| GenBank Accession | Study | Location | SNP Genotype |  |  |
| --- | --- | --- | --- | --- | --- |
|  |  |  | 15564 (A/G) | 15760 (C/T) | 15779 (C/T) |
| KJ522887.1 | McLoad 2014 | West | A | T | C |
| KJ522888.1 | McLoad 2014 | West | A | T | C |
| KJ522889.1 | McLoad 2014 | West | A | T | C |
| KJ522890.1 | McLoad 2014 | West | G | T | T |
| KJ522891.1 | McLoad 2014 | West | A | T | C |
| KJ522892.1 | McLoad 2014 | West | A | T | C |
| KJ522893.1 | McLoad 2014 | West | G | T | T |
| KJ522894.1 | McLoad 2014 | West | A | T | C |
| KJ522895.1 | McLoad 2014 | West | G | T | C |
| **KJ522896.1** | **McLoad 2014** | **West** | **A** | **C** | **C** |
| KJ522897.1 | McLoad 2014 | West | A | T | C |
| **KJ522898.1** | **McLoad 2014** | **West** | **A** | **C** | **C** |
| KJ522899.1 | McLoad 2014 | West | G | T | T |
| KJ522900.1 | McLoad 2014 | West | G | T | T |
| KJ522901.1 | McLoad 2014 | West | G | T | T |
| **KJ522902.1** | **McLoad 2014** | **West** | **A** | **C** | **C** |
| **KJ522903.1** | **McLoad 2014** | **West** | **A** | **C** | **C** |
| **KJ522904.1** | **McLoad 2014** | **West** | **A** | **C** | **C** |
| KJ522905.1 | McLoad 2014 | West | A | T | C |
| KJ522906.1 | McLoad 2014 | West | G | T | C |
| KJ522907.1 | McLoad 2014 | West | A | T | C |
| KJ522908.1 | McLoad 2014 | West | A | T | C |
| KJ522909.1 | McLoad 2014 | West | A | T | C |
| KJ522910.1 | McLoad 2014 | West | G | T | T |
| **KJ522911.1** | **McLoad 2014** | **West** | **A** | **C** | **C** |
| KJ522912.1 | McLoad 2014 | West | G | T | C |
| KJ522913.1 | McLoad 2014 | West | A | T | C |
| KJ522914.1 | McLoad 2014 | West | A | T | C |
| KJ522915.1 | McLoad 2014 | West | A | T | C |
| KJ522916.1 | McLoad 2014 | West | A | T | C |
| KJ522917.1 | McLoad 2014 | West | A | T | C |
| KJ522918.1 | McLoad 2014 | West | A | T | C |
| KJ522919.1 | McLoad 2014 | West | A | T | C |
| **KJ522920.1** | **McLoad 2014** | **West** | **A** | **C** | **C** |
| **KJ522921.1** | **McLoad 2014** | **West** | **A** | **C** | **C** |
| **KJ522922.1** | **McLoad 2014** | **West** | **A** | **C** | **C** |
| MF166700.1 | Andersen 2017 | East | G | T | C |
| MF166701.1 | Andersen 2017 | East | A | T | T |
| MF166702.1 | Andersen 2017 | East | G | T | C |
| MF166703.1 | Andersen 2017 | East | G | T | C |
| MF166704.1 | Andersen 2017 | East | A | T | T |
| MF166705.1 | Andersen 2017 | East | G | T | T |
| MF166706.1 | Andersen 2017 | East | G | T | T |
| MF166707.1 | Andersen 2017 | East | A | T | C |
| MF166708.1 | Andersen 2017 | East | A | C | T |
| MF166709.1 | Andersen 2017 | East | G | T | C |
| MF166710.1 | Andersen 2017 | East | G | T | T |
| MF166711.1 | Andersen 2017 | East | G | T | C |
| MF166712.1 | Andersen 2017 | East | A | C | T |
| MF166713.1 | Andersen 2017 | East | A | T | C |
| MF166714.1 | Andersen 2017 | East | A | T | T |
| MF166715.1 | Andersen 2017 | East | G | T | C |
| MF166716.1 | Andersen 2017 | East | G | T | C |
| MF166717.1 | Andersen 2017 | East | G | T | T |
| MF166718.1 | Andersen 2017 | East | A | T | T |
| MF166719.1 | Andersen 2017 | East | G | T | T |
| MF166720.1 | Andersen 2017 | East | A | T | T |
| MF166721.1 | Andersen 2017 | East | A | T | T |
| MF166722.1 | Andersen 2017 | East | G | T | T |
| MF166723.1 | Andersen 2017 | East | A | T | T |
| MF166724.1 | Andersen 2017 | East | A | T | T |
| KU710183.1 | Bachman 2016 | East | A | T | T |
| KU710184.1 | Bachman 2016 | East | G | T | T |
| KU710185.1 | Bachman 2016 | East | G | T | C |
| KU710186.1 | Bachman 2016 | East | A | T | T |
| KU710187.1 | Bachman 2016 | East | G | T | C |
| KU710189.1 | Bachman 2016 | East | A | T | T |
| KU710190.1 | Bachman 2016 | East | A | T | T |
| KU710191.1 | Bachman 2016 | East | A | T | T |
| KU710192.1 | Bachman 2016 | East | G | T | T |
| KU710193.1 | Bachman 2016 | East | G | T | C |
| KU710194.1 | Bachman 2016 | East | G | T | T |
| KU710195.1 | Bachman 2016 | East | G | T | T |
| KU710196.1 | Bachman 2016 | East | G | T | T |
| KU710197.1 | Bachman 2016 | East | A | T | T |
| KU710198.1 | Bachman 2016 | East | A | T | T |
| KU710199.1 | Bachman 2016 | East | G | T | C |
| KU710200.1 | Bachman 2016 | East | G | T | T |
| EU728544.1 | Lindquist 2009 | East | A | T | T |
| EU728545.1 | Lindquist 2009 | East | A | T | T |
| EU728546.1 | Lindquist 2009 | East | A | T | T |
| EU728547.1 | Lindquist 2009 | East | A | T | C |
| EU728548.1 | Lindquist 2009 | East | A | T | T |
| **EU728549.1** | **Lindquist 2009** | **West** | **A** | **C** | **C** |
| **EU728550.1** | **Lindquist 2009** | **West** | **A** | **C** | **C** |
| **EU728551.1** | **Lindquist 2009** | **West** | **A** | **C** | **C** |
| **EU728552.1** | **Lindquist 2009** | **West** | **A** | **C** | **C** |
| **EU728553.1** | **Lindquist 2009** | **West** | **A** | **C** | **C** |
| EU728554.1 | Lindquist 2009 | East | A | T | T |
| EU728555.1 | Lindquist 2009 | East | A | T | T |
| EU728556.1 | Lindquist 2009 | East | A | T | T |
| EU728557.1 | Lindquist 2009 | East | A | T | T |
| EU728558.1 | Lindquist 2009 | East | A | T | T |
| **EU728559.1** | **Lindquist 2009** | **West** | **A** | **C** | **C** |
| **EU728560.1** | **Lindquist 2009** | **West** | **A** | **C** | **C** |
| **EU728561.1** | **Lindquist 2009** | **West** | **A** | **C** | **C** |
| EU728565.1 | Lindquist 2009 | East | A | C | T |
| EU728566.1 | Lindquist 2009 | East | A | T | T |
| EU728567.1 | Lindquist 2009 | East | A | T | T |
| EU728568.1 | Lindquist 2009 | East | A | T | C |
| EU728569.1 | Lindquist 2009 | East | G | T | T |
| EU728570.1 | Lindquist 2009 | East | G | C | T |
| EU728571.1 | Lindquist 2009 | East | G | T | C |
| EU728572.1 | Lindquist 2009 | East | G | T | C |
| EU728573.1 | Lindquist 2009 | East | G | T | C |

## Supplemenatary Figures

**Figure S1.**
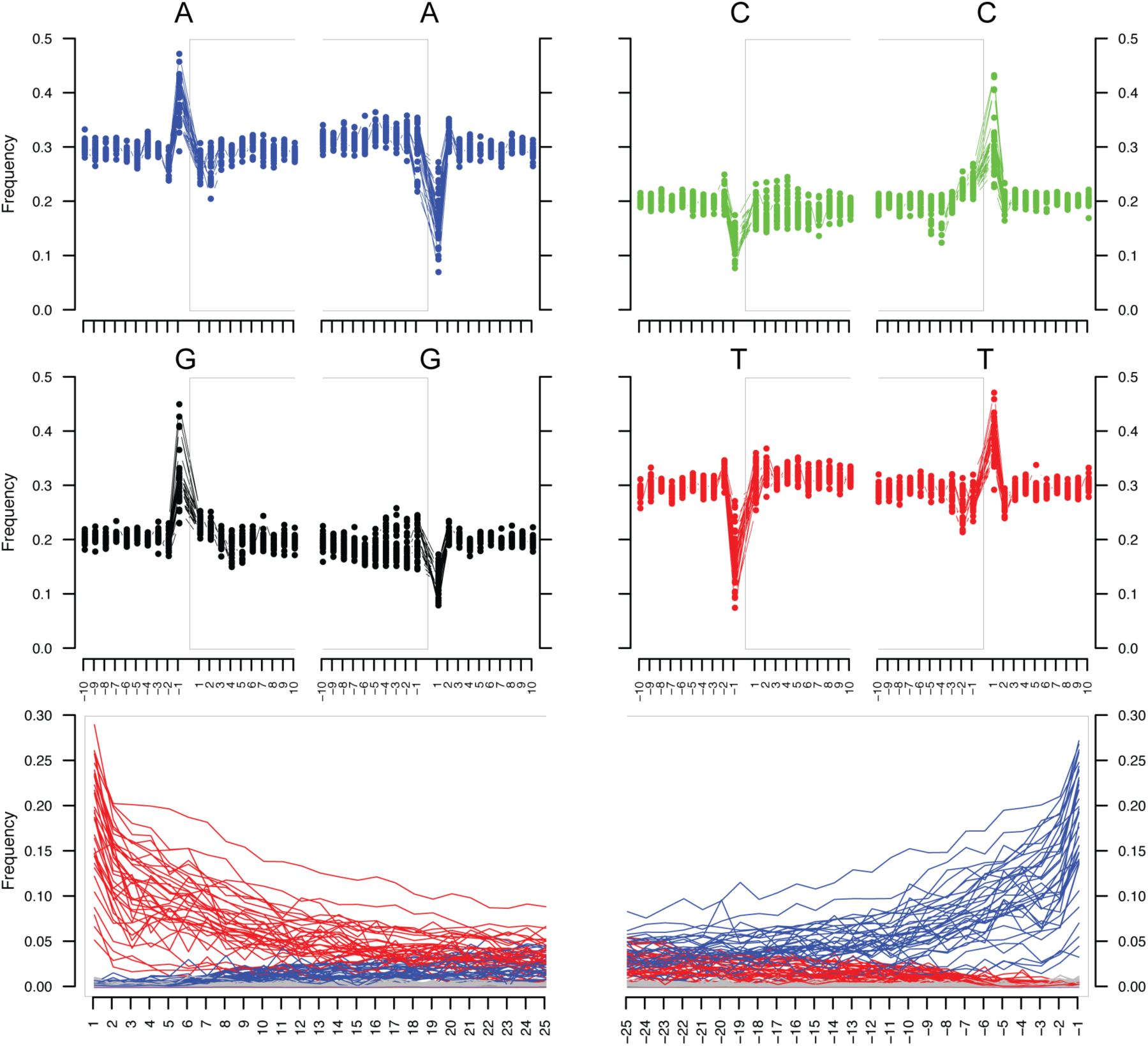
aDNA fragmentation and mis-incorporation patterns of sequencing read data from 37 archaeological Atlantic walrus samples. All samples show the typical fragmentation (top four panels), elevated 5’-end C->T (bottom left panel) and elevated 3’-end G->A substitution patterns (bottom right panel) expected from sequencing authentic aDNA data. Patterns were obtained using MapDamage v. 2.0.6 after down-sampling BAM files to 1,000,000 reads if applicable.

**Figure S2.**
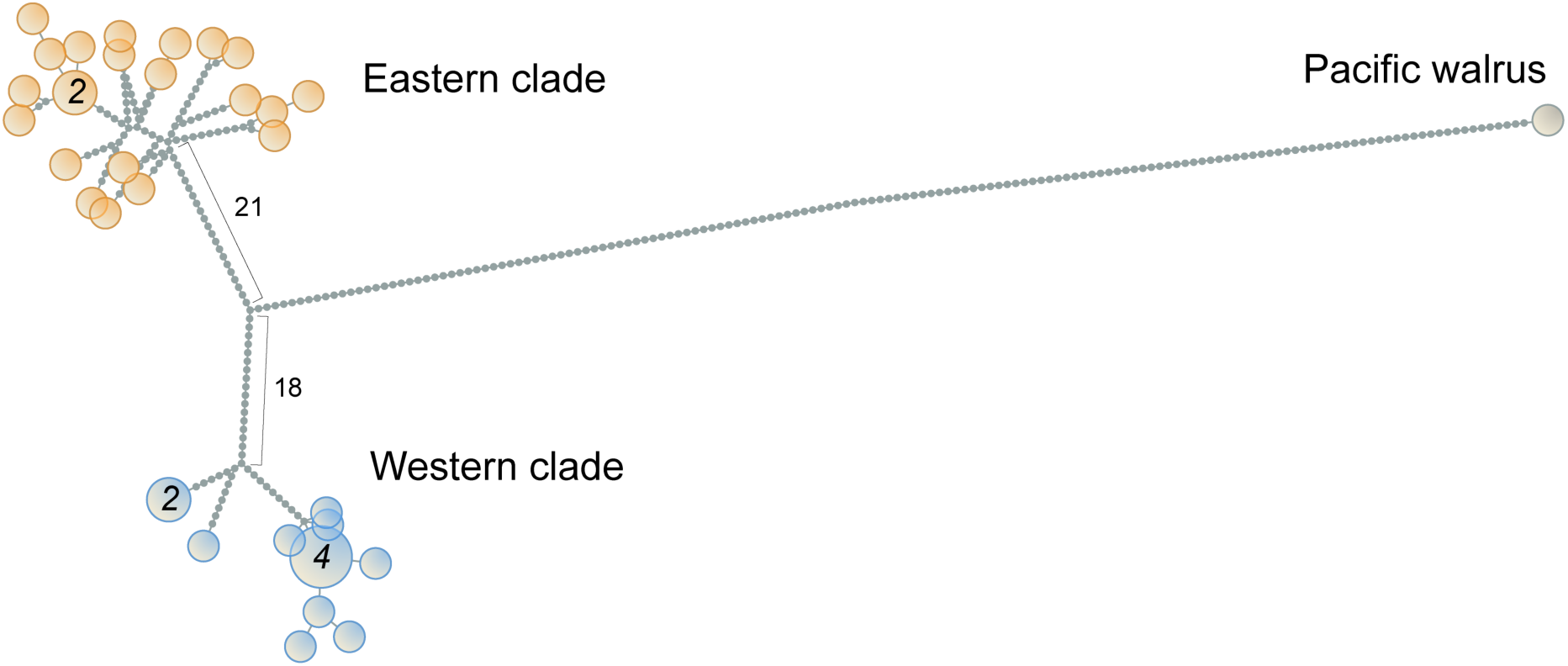
Haplotype genealogy graph of 37 archaeological Atlantic walruses and one Pacific walrus. Haplotypes belonging to the Western clade (*blue*), Eastern clade (*orange*) or the Pacific walrus (*grey*) are separated by a number of substitutions (grey edges indicated with a number for the two main Atlantic walrus branches). Circle size reflects the number of specimens with an identical haplotype, and where this is >1 the number is specified within the circle.

**Figure S3.**
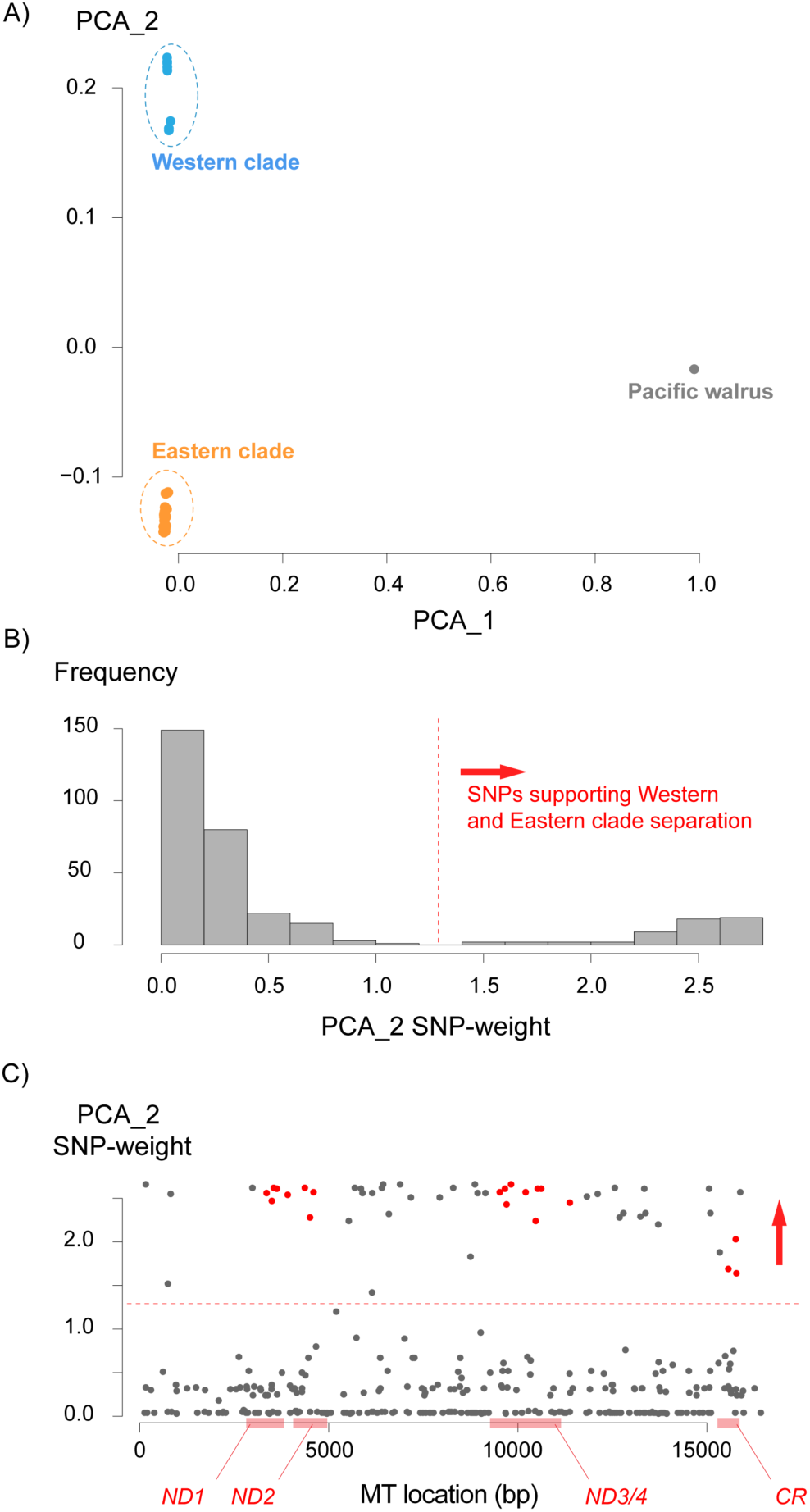
Genetic population structure based on whole mitochondrial genome data in 37 archaeological Atlantic walruses and one Pacific walrus. **a)** Principle component analysis (PCA) based on 346 SNPs using smartPCA, EIGENSOFT v.6.1.4. The first principle component significantly (eigenvalue = 15.5, Tracy-Widom (TW) stat = 1.76, *p*-value = 0.01) differentiates the Pacific walrus (*grey*) from the Atlantic specimens. The second principle component significantly (eigenvalue = 7.01, TW stat = 4.26, *p*-value = 0.0001) differentiates the western (*blue*) and eastern (*orange*) Atlantic walrus clades. **b)** A bimodal distribution characterizes the SNP-weightings of the second Principle Component. SNPs with a weighting above 1.25 support the western and eastern differentiation, and those with the highest values are exclusively associated with either clade. **c)** SNPs supporting the western and eastern differentiation are located throughout the mitochondrial genome with a subset of SNPs (*red*) located in those regions (*ND1*, *ND2*, *ND3*/*4* and the control region *CR, red italic*) investigated in previous studies.

